# Marine heatwaves depress metabolic activity and impair cellular acid-base homeostasis in reef-building corals regardless of bleaching susceptibility

**DOI:** 10.1101/2021.02.23.432550

**Authors:** Teegan Innis, Luella Allen-Waller, Kristen Taylor Brown, Wesley Sparagon, Christopher Carlson, Elisa Kruse, Ariana S. Huffmyer, Craig E. Nelson, Hollie M. Putnam, Katie L. Barott

## Abstract

Ocean warming is causing global coral bleaching events to increase in frequency, resulting in widespread coral mortality and disrupting the function of coral reef ecosystems. However, even during mass bleaching events, many corals resist bleaching despite exposure to abnormally high temperatures. While the physiological effects of bleaching have been well documented, the consequences of heat stress for bleaching resistant individuals are not well understood. In addition, much remains to be learned about how heat stress affects cellular level processes that may be overlooked at the organismal level, yet are crucial for coral performance in the short term and ecological success over the long term. Here we compared the physiological and cellular responses of bleaching resistant and bleaching susceptible corals throughout the 2019 marine heatwave in Hawai‘i, a repeat bleaching event that occurred four years after the previous regional event. Relative bleaching susceptibility within species was consistent between the two bleaching events, yet corals of both resistant and susceptible phenotypes exhibited pronounced metabolic depression during the heatwave. At the cellular level, bleaching susceptible corals had lower intracellular pH than bleaching resistant corals at the peak of bleaching for both symbiont-hosting and symbiont-free cells, indicating greater disruption of acid-base homeostasis in bleaching susceptible individuals. Notably, cells from both phenotypes were unable to compensate for experimentally induced cellular acidosis, indicating that acid-base regulation was significantly impaired at the cellular level even in bleaching resistant corals and in cells containing symbionts. Thermal disturbances may thus have substantial ecological consequences, as even small reallocations in energy budgets to maintain homeostasis during stress can negatively affect fitness. These results suggest concern is warranted for corals coping with ocean acidification alongside ocean warming, as the feedback between temperature stress and acid-base regulation may further exacerbate the physiological effects of climate change.

## Introduction

Climate change has led to catastrophic losses in live coral cover on reefs around the world as ocean warming has intensified and marine heatwaves have increased in frequency and severity (T. P. Hughes, Anderson, et al. 2018; Hoegh-Guldberg et al. 2007). These anomalous thermal stress events can trigger coral bleaching, the loss of photosynthetic algal endosymbionts (family Symbiodiniaceae) from their animal host (Weis 2008; Oakley and Davy 2018). This loss of the host’s primary food source creates an energetic deficit that often results in mortality (McClanahan 2004; T. P. Hughes, Anderson, et al. 2018). Mass mortality events associated with bleaching have led to drastic declines in coral cover on reefs globally, altering the composition of the entire reef community and disrupting reef function (T. P. Hughes, Kerry, et al. 2018; Leggat et al. 2019; Fordyce et al. 2019). Recovery from bleaching is possible, however, and some corals can compensate for loss of the autotrophic contribution of their algal symbionts by increasing heterotrophic feeding and catabolizing endogenous reserves (Grottoli, Rodrigues, and Palardy 2006; A. D. Hughes and Grottoli 2013; Tolosa et al. 2011). Despite this flexibility, as the frequency of coral bleaching increases, the shortened duration between events may be insufficient for complete recovery of many survivors (Grottoli et al. 2014; Sale et al. 2019; Schoepf et al. 2015). These energetic or functional deficits during recovery from bleaching can increase coral susceptibility to subsequent stressors, including but not limited to temperature, disease, competition, and ocean acidification (Ward, Harrison, and Hoegh-Guldberg 2000; Muller, Bartels, and Baums 2018; Brown et al. 2019). Short intervals between heatwaves can also change the relative performance between coral species, as differential investments in resistance versus recovery strategies (e.g. (Matsuda et al. 2020)) may enable some species to withstand single but not repeat bleaching events (Grottoli et al. 2014).

Understanding the diversity of coral heat stress responses both within and between species is critical for predicting future coral performance in a changing ocean. While coral bleaching is a dramatic and easily observable heat stress response (Putnam et al. 2017; Bollati et al. 2020), elevated temperatures can also trigger less visible stress responses in the animal host, even if the symbiosis remains intact (Rädecker et al. 2021). All animals have a range of temperatures to which they are adapted, and once exposed to temperatures beyond this optimal range, can activate a suite of heat stress responses and divert energetic investment towards maintenance processes like homeostasis, at the expense of other more immediately expendable processes like growth and reproduction (H. Pörtner 2008). Scleractinian corals live near the upper edge of their thermal optimum, and even small increases above their typical maximum summer temperatures can activate molecular heat stress response pathways (Cleves et al. 2020; Seneca and Palumbi 2015; Bay and Palumbi 2015). At the organismal level, heat stress can also lead to metabolic depression in corals even when the symbiosis remains intact (Edmunds, Cumbo, and Fan 2011; Bernardet et al. 2019; Silbiger et al. 2019). While this metabolic depression can be important for promoting survival during stress in the short term (H. Pörtner 2008), it can have long-term negative consequences for energetic status (Grottoli, Rodrigues, and Palardy 2006) and ecological success (Anthony et al. 2009). For example, in corals, this reallocation of energy use and decreased metabolism during heat stress can lead to reductions in feeding (Ferrier-Pagès et al. 2010), calcification (Cantin and Lough 2014), and reproduction (Levitan et al. 2014; Fisch et al. 2019; Baird and Marshall 2002), all critical processes for coral survival. Furthermore, bleaching exacerbates these energetic deficits, causing more pronounced effects on organismal performance, with bleaching susceptible individuals suffering greater reductions in growth, disease resistance and reproduction in the months and years following heat stress than bleaching resistant conspecifics (Levitan et al. 2014; Fisch et al. 2019; Matsuda et al. 2020). Therefore, it is important to understand the influence of marine heatwaves on coral physiology regardless of bleaching susceptibility, because even bleaching resistant corals may suffer energetic losses, ultimately decreasing coral fitness and overall reef function.

Declines in metabolic performance due to heat stress can also trigger changes in cellular processes such as acid-base regulation of intracellular and external body fluids (H. Pörtner 2008). Maintaining a steady pH is crucial for all living organisms, as nearly all metabolic processes require a narrow pH range in order to function (Tresguerres et al. 2020). Thus, characterizing the influence of temperature on acid-base homeostasis mechanisms is essential for understanding coral physiological responses to a changing climate. In other marine invertebrates, heat stress can cause declines in intracellular pH (pH_i_) (i.e. the homeostatic setpoint) and a decreased ability to maintain the pH setpoints of extracellular body fluids, generally a result of reduced metabolism, protein synthesis and ion exchange rates (H. Pörtner 2008). Understanding the physiological interactions between temperature stress and acid-base homeostasis is critical for predicting coral performance and acclimatization potential in a changing environment.

However, little is known about how climate change affects acid-base homeostasis in corals. The potential effects of heat stress on acid-base regulation pose a particular challenge for maintaining coral calcification, as biomineralization is highly pH-dependent (Barott, Venn, et al. 2020; Venn et al. 2011; McCulloch et al. 2012). Indeed, recent studies have found that temperature stress alters the pH of the external calcifying fluid and depresses calcification (Schoepf et al. 2021; Guillermic et al. 2021), highlighting the need to better understand the physiology of coral climate change responses across biological scales.

In order to assess how marine heatwaves influence the interactions between coral metabolism, symbiosis and cellular acid-base homeostasis, we characterized the physiological responses of bleaching resistant and bleaching susceptible individuals of two dominant reef-building corals in Hawai’i, *Montipora capitata* and *Porites compressa*, to a repeat bleaching event that occurred in Kāne’ohe Bay, O’ahu in 2019. Coral bleaching susceptibility of these individuals was initially quantified during the previous bleaching event in 2015, when they were tagged as a resource for future studies (Matsuda et al. 2020). During the subsequent 2019 heatwave, a suite of physiological variables was quantified at both the cellular and organismal level every two weeks throughout the peak of the heatwave and the responses of each species and phenotype were compared. This study takes an important step towards understanding the effects of repeat heatwaves on coral physiology across biological scales and helps disentangle the effects of heat stress versus bleaching on coral physiology by comparing responses between conspecific individuals with contrasting bleaching susceptibilities.

## Materials and Methods

### Study site

Coral monitoring and collections took place in Kāne’ohe Bay, O’ahu, Hawai’i. Kāne’ohe Bay is a lagoon that harbors a system of coral-dominated patch and fringing reefs protected by a barrier reef which restricts circulation and results in temperatures 1-2°C higher than the surrounding ocean in the summer months (Bahr, Jokiel, and Rodgers 2015), although coral bleaching has only been observed during anomalous heatwaves (Bahr, Rodgers, and Jokiel 2017). This study was conducted at a coral-dominated patch reef (PR) in the outer lagoon (PR13; 21.4515°N, 157.7966°W), where the most abundant reef-building corals were *Montipora capitata* and *Porites compressa*. This region of Kāne’ohe Bay is characterized by relatively short seawater residence times (<24 hours, Lowe et al. 2009) and diel pH ranges 2 – 3 fold higher than other regions of the bay with longer residence times (Barott, Huffmyer, et al. 2020; Page et al. 2018). Hourly sea surface temperatures on the reef from the 2015 and 2019 heatwaves were obtained from the NOAA temperature sensor on Moku o Lo’e in Kāne’ohe Bay (Station ID: 1612480). Cumulative heat stress in the form of degree heating weeks (DHW) was calculated from 1 May to 31 December as the hourly accumulation of sea surface temperatures over the previous 12 weeks above the bleaching threshold (29°C), determined as 1°C above the local maximum monthly mean (MMM, 28.0°C; Jury and Toonen 2019; Fig. 1A).

**Figure 1.**
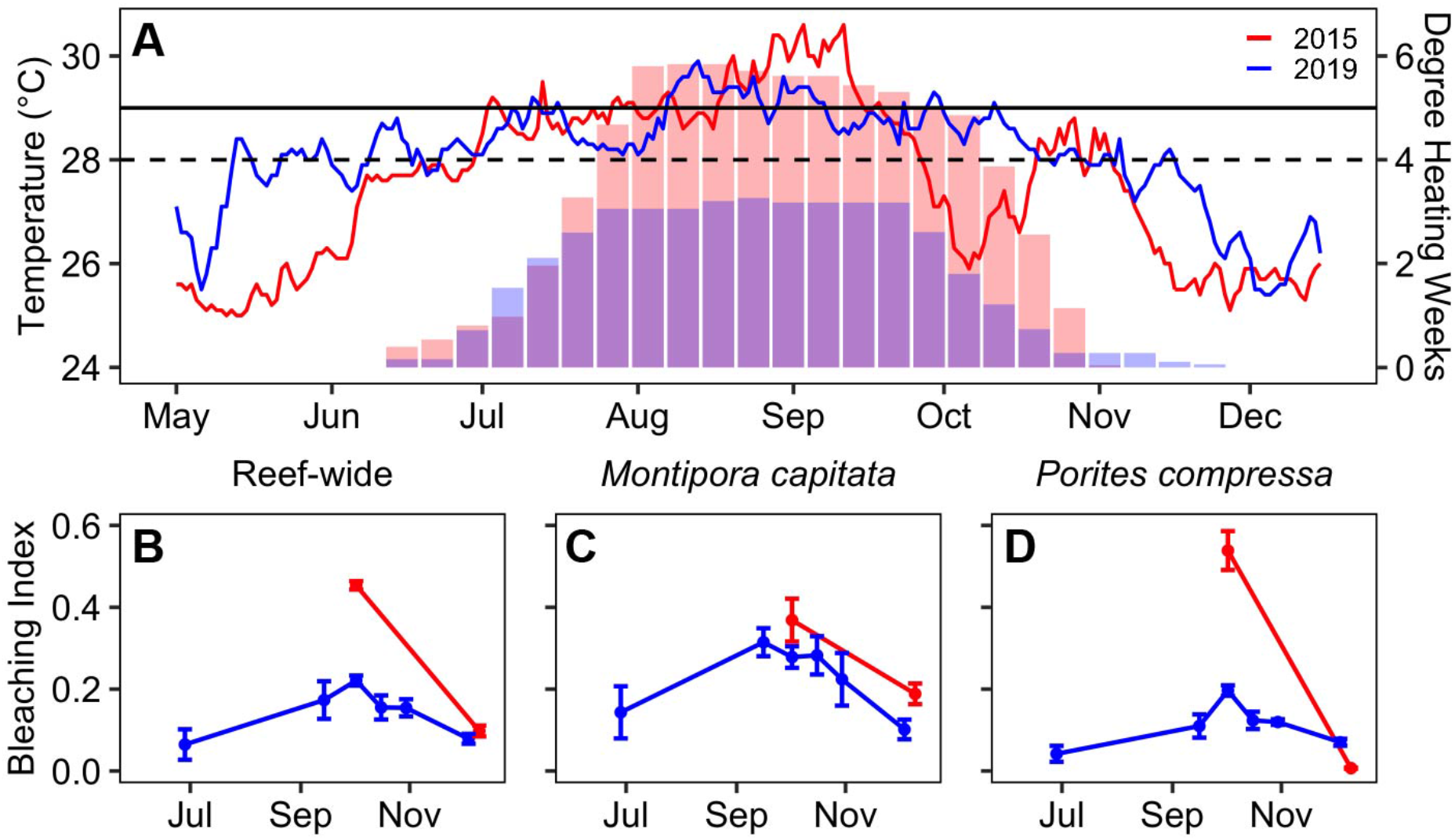
Repeat marine heatwaves in 2015 and 2019 led to coral bleaching in Kāne‘ohe Bay, O‘ahu, Hawai‘i. A) Maximum daily temperatures (lines) and accumulated heat stress (DHW; bars) at the Moku o Lo‘e (Coconut Island) monitoring station. Dashed horizontal line indicates local maximum monthly mean (MMM). Solid horizontal line indicates local coral bleaching threshold (MMM+1°C). Bleaching index was calculated from benthic surveys (n=4; mean ± SE) at the B) reef-wide scale and for species of interest, C) *Montipora capitata* and D) *Porites compressa*.

### Monitoring of the extent and severity of coral bleaching

Benthic community composition and the extent and severity of coral bleaching was quantified before the heatwave (28 June 2019), every two weeks during thermal stress (16 September – 30 October 2019) and following the heatwave (4 December 2019). At each time point, 0.33 m^2^ quadrats were photographed on the benthos at 2 m intervals along a 40 m transect laid parallel to the reef crest, totaling 20 images per transect. Transects were conducted at both 1 m and 3 m depths (n=2 per depth), for a total of four transects per time point. Images were taken using an underwater camera (Olympus TG-5) with size and color standards. The extent of coral bleaching was determined using CoralNet (Beijbom et al. 2015), with 50 random points per image chosen for benthic identification. Corals were identified to species and categorized as: (1) pigmented/not bleached, (2) pale/moderately bleached, or (3) white/severely bleached. The bleaching index (BI), a measure of the extent of bleaching at the reef level, was calculated as the weighted average of the percentage (*P*) of observations in each bleaching category (1-3) with the equation: *BI = (0*P*_*1*_ *+ 1*P*_*2*_ *+ 2*P*_*3*_*)/2*, as in (McClanahan 2004).

### Identification and collection of bleaching susceptible and bleaching resistant corals

During the peak of the 2015 bleaching event, 10 pairs of severely bleached and fully pigmented individuals of *M. capitata* and *P. compressa* adjacent to each other on the reef (i.e., each pair consisted of one bleached and one pigmented colony) were identified and tagged (Fig. 2A,D; Matsuda et al. 2020). These pairs were photographed with size and color standards before the marine heatwave (19 July 2019), every two weeks during thermal stress (16 September – 30 October 2019) and following the heatwave (24 January 2020). Bleaching severity, a measure of individual colony bleaching, was visually determined for each colony as follows: (1) 0% bleached, (2) <20% bleached, (3) 20-50% bleached, (4) 50-80% bleached and (5) >80% bleached, as in (McClanahan 2004). For each of the four time points between September and October, one 4–5 cm fragment from each colony was also sampled. Fragments were held in individual bags of ambient seawater during transit to the Hawai’i Institute of Marine Biology (HIMB), transferred to an outdoor flow-through seawater table (ambient seawater sourced from Kāne’ohe Bay) within three hours of collection, and held there for a maximum of three days until assessment of *in vivo* coral performance.

**Figure 2.**
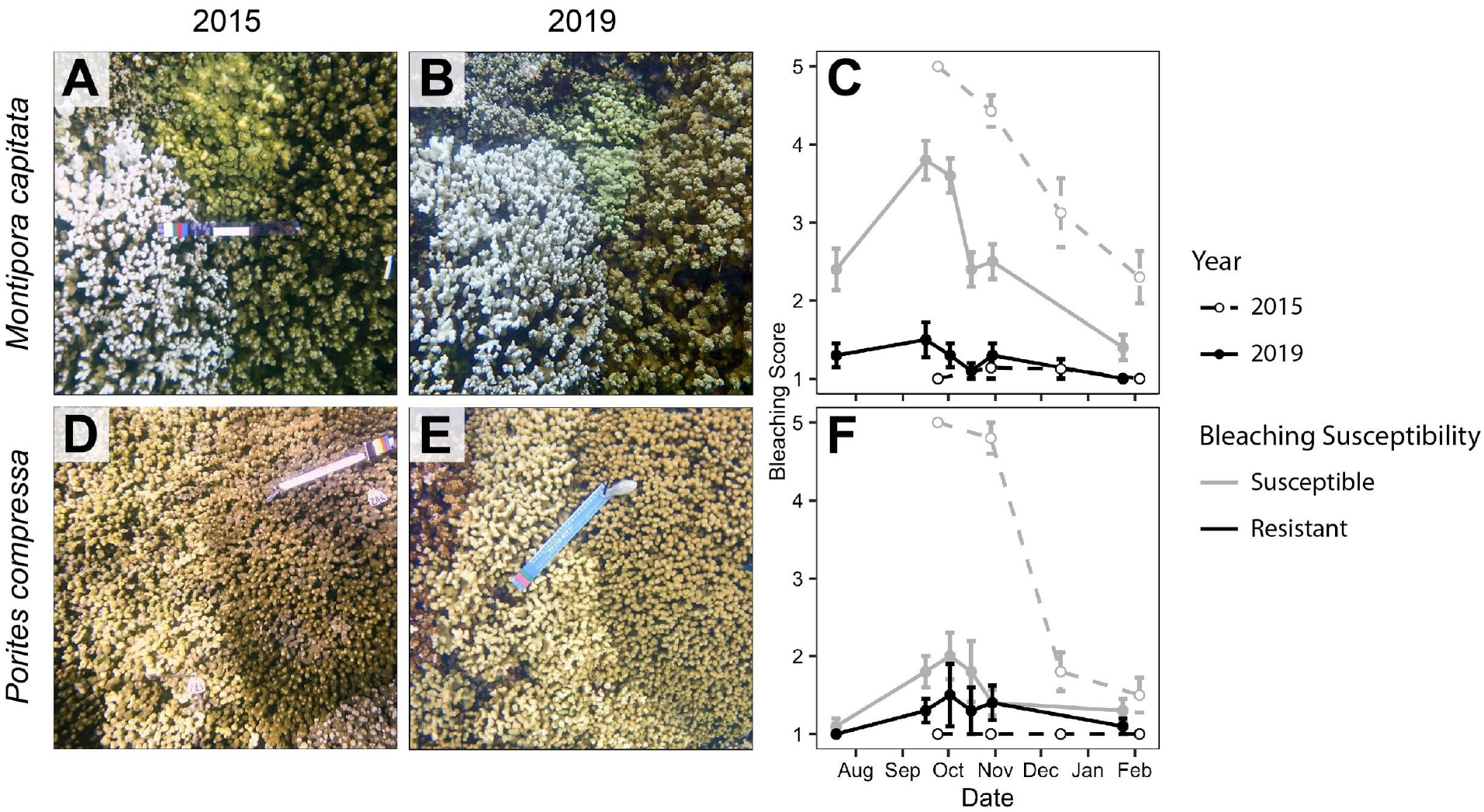
Response of bleaching resistant and bleaching susceptible corals to marine heatwaves in 2015 and 2019. Representative images of an *Montipora capitata* pair during the A) 2015 and B) 2019 heatwaves. C) Mean bleaching scores of bleaching susceptible and bleaching resistant *M. capitata* throughout each heatwave (N=10; mean ± SEM). Representative images of a *Porites compressa* pair during the D) 2015 and E) 2019 heatwaves. F) Mean bleaching scores of bleaching susceptible and bleaching resistant *P. compressa* throughout each heatwave (N=10; mean ± SEM). Dec 2015 - Feb 2019, N=10 pairs; Sept-Oct 2015, N=3-7 pairs.

### In vivo coral performance

Dark-adapted photochemical yield (F_v_/F_m_) was measured on each coral fragment using a Diving-PAM (Walz GmbH, Germany) approximately 1 hour after sunset on the same day of collection. Measurements were made using the Diving-PAM 5-mm diameter fiber-optic probe at a standardized 5 mm above the coral tissue after F_0_ stabilized. The Diving-PAM settings were set to a measuring light intensity of 5, gain of 2, and saturation pulse intensity of 5. Within 3 days of collection, rates of photosynthesis and light enhanced dark respiration (LEDR) were subsequently measured on each coral fragment by quantifying oxygen evolution and consumption as follows. Each coral fragment was placed in a 250 mL sealed chamber filled with ambient seawater surrounded by a temperature-controlled water jacket to maintain a constant temperature (ambient: 25-27°C). Seawater in the chambers was constantly mixed using a magnetic stir bar. Temperature and dissolved oxygen concentrations were measured using a Pt100 temperature probe and PSt7 oxygen optode (PreSens), respectively, inserted through a port in the lid of each chamber. Oxygen optodes were calibrated on each measurement day with a 0% oxygen solution (0.01 g mL^-1^ NaSO_3_) and 100% air saturated seawater. Oxygen evolution rates were measured at steady increments of light (112–726 µmol m^-2^ s^-1^, EcoTech Radion XR30w Pro), increasing light intensity only after a steady slope was achieved for all fragments for at least 10 min. After the maximum light level, the lights were turned off and oxygen consumption rates were measured until a steady slope was achieved for at least 10 minutes to use for LEDR calculations. Temperature and oxygen data were recorded every 3 seconds by an OXY-10 ST (PreSens). A control chamber (seawater only) was analyzed in parallel for each run. Corals were then snap frozen and stored at -80°C until further processing.

Photosynthetic and respiration rates were calculated from volumetric oxygen production and consumption rates (i.e., µmol O_2_ L^-1^ min^-1^) by multiplying the oxygen concentration changes and volume of water in each chamber (chamber volume – coral volume, L) and accounting for background oxygen flux rates by subtracting the rate of the corresponding seawater-only control chamber. The volume of each coral fragment was measured via the water displacement method. Photosynthesis-irradiance (PI) curves (Fig. S1) were generated by curve fitting to the Platt model (Platt, Gallegos, and Harrison 1980) in order to extract alpha, Ik, and Pmax. LEDR was calculated from the dark period following the PI curve maximum step. Metabolic rates were normalized to surface area (described below) and gross photosynthetic rates were determined by subtracting the oxygen consumption rates (LEDR) from oxygen production for each fragment.

### Physiological analyses

Tissue was removed from each fragment using an airbrush containing phosphate-buffered saline (PBS) solution. The resulting tissue slurry was homogenized at 25,000 rpm for 10 seconds using a tissue homogenizer (Thermo Fisher Scientific) and aliquoted for subsequent assays. For symbiont cell counts, tissue homogenate was homogenized at 25,000 rpm for 10 seconds followed by needle shearing with a 22-gauge needle. Algal cells were then pelleted by centrifugation at 7,000 g for 5 min and resuspended in 0.1% sodium dodecyl sulfate (SDS) in 0.22 µm filtered seawater (FSW). Symbiodiniaceae concentrations were determined following (Krediet et al. 2015) using a Millipore Guava flow-cytometer (Guava easyCyte 5HT) and modifications to the methods outlined here. Three technical replicates were quantified for each sample. Symbiodiniaceae cells were excited with a blue laser (488 nm) and identified by analyzing forward scatter and red autofluorescence in GuavaSoft 3.4 with the same gating for all samples. Chlorophyll was extracted in 100% acetone. Briefly, tissue homogenate was centrifuged at 14,000 rpm for 3 min at 4°C and the supernatant was removed. The remaining pellet was incubated in 100% acetone for 32-48 hr in the dark at -20°C. The samples were then centrifuged at 14,000 rpm for 3 min at 4°C. The supernatant was transferred to a 96-well flat bottom glass plate and absorbance was quantified in duplicate. The concentrations of chlorophyll a and c_2_ were determined by measuring absorbance at 630 nm, 663 nm and 750 nm on a plate reader (BioTek PowerWave XS2) and calculated from the equations in (Jeffrey and Humphrey 1975) for dinoflagellates in 100% acetone, correcting for the 0.6 cm path length of the 96-well quartz plate: *Chl a = (11*.*43(A*_*663*_ *- A*_*750*_*/PL) – 0*.*64(A*_*630*_ *– A*_*750*_*/PL))/(mL homogenate); Chl c2 = (27*.*09 (A*_*630*_ *- A*_*750*_*/PL) – 3*.*63(A*_*663*_ *- A*_*750*_*/PL))/(mL homogenate)*.

Soluble protein content was analyzed in triplicate via the Bradford method using Coomassie Plus Bradford assay reagent (Pierce Biotechnology). The crude tissue homogenate was analyzed to obtain a measure of holobiont protein. For the host fraction, symbionts were removed from the crude homogenate by centrifugation at 10,000 g for 10 min at 4°C, and the resulting supernatant was analyzed. The samples were mixed with the reagent on a plate shaker for 30 sec, incubated for 10 min at room temperature and again mixed on a plate shaker for 30 sec. The sample absorbance was then measured at 595 nm on a plate reader (BioTek ELx808). Protein concentration was calculated against a bovine serum albumin (BSA) standard curve run alongside the samples and normalized to skeletal surface area. Tissue biomass was determined here as ash-free dry weight (AFDW). A known volume of each homogenate was dried at 60°C for 24 hr until a constant weight was achieved. After the dry weight was recorded, the homogenates were then burned in a muffle furnace at 450°C for 6 hr. The samples were allowed to cool in the furnace before being weighed and the ash weight recorded. The difference between the ash weight and the dry weight was calculated to determine the AFDW of each fragment.

Total antioxidant capacity was measured using the OxiSelect Total Antioxidant Capacity (TAC) Assay Kit (Cell Biolabs). The tissue homogenate was first centrifuged at 10,000 g for 10 min at 4°C prior to loading on the 96-well plate. Each sample and uric acid standard was run in two technical replicates per manufacturer’s instructions. Net maximum absorbance values were measured at 490 nm on a plate reader (BioTek ELx808) at the initial and final time points and the difference of those was used to calculate TAC in µM against the uric acid standard curve. Uric acid equivalents were converted to copper reducing equivalents (CRE) per manufacturer’s instructions and normalized to host protein.

After tissue was removed, skeletons were soaked in 10% bleach for approx. 12 h and then dried at 60°C for approx. 12 h until a constant weight was reached. Surface area was determined by the single wax dipping method (Veal et al. 2010). Each skeleton fragment was pre-weighed before being dipped in paraffin wax, after which the final weight was recorded. The change in weight due to wax addition was compared against a standard curve of dipped wooden dowels of known surface area to calculate the skeletal surface area of each fragment.

### Assessment of coral intracellular acid-base homeostasis

Intracellular pH (pH_i_) of *M. capitata* was measured at the peak of the bleaching response (28 September – 2 October 2019) using established methods (Barott et al. 2017). Briefly, coral fragments were held in the dark for 30 min, then cells were isolated and loaded with 10 µM SNARF-1 AM, 0.1% DMSO, and 0.01% Pluronic F-27 in filtered seawater (FSW) for 30 min at 25°C in the dark. Cells were then pelleted and resuspended in either ambient FSW (pH 8.0) to measure the pH_i_ setpoint or HCl-acidified FSW (pH 7.4) to measure their response to acidosis. Seawater pH was confirmed immediately prior to resuspension (accupHast pH probe and accumet pH meter, Fisherbrand) and readjusted if needed. All cells were imaged in a glass-bottomed dish using an inverted confocal microscope (Zeiss LSM 710) at 63X magnification (Plan-Apochromat Oil DIC M27 objective, numerical aperture = 1.4). All samples were excited at 561 nm (DPSS, 2% power, 458/561 main beam splitter). SNARF-1 fluorescence emission was acquired in two channels simultaneously at 585 and 640 +/- 10 nm, using PMT detectors (gain = 500). Images were acquired at 12 bits/pixel using a scan speed of 261.8 Hz. The microscope stage was maintained at 25.0°C. A total of 8–10 cells containing algal symbionts (symbiocytes) and 8–10 cells without symbionts (non-symbiocytes) were imaged from each coral. To measure acid stress response, SNARF-1-loaded cells from the same fragment were resuspended in HCl-acidified FSW (pH 7.4), and imaged over time (5, 20, 30, 50, and 70 min after acid stress). A total of 5–10 cells of each cell type were measured per time point. SNARF-1 fluorescence emission was quantified in ImageJ by drawing two regions of interest (ROI) per cell within the coral cytoplasm. A region drawn in the surrounding medium was used to subtract background fluorescence, and each region of interest (ROI) fluorescence ratio was calculated and converted to pH using a calibration curve generated as previously described (Venn *et al*., 2009). Cytosolic acidification magnitude (i.e. acidosis) was calculated for each coral as the difference between initial pH_i_ and pH_i_ after 5 minutes of acidification. The pH_i_ recovery rate was calculated for each coral by regressing pH_i_ measurements of each cell against their corresponding image timestamps for all times after the first 5 min. Attempts to measure pH_i_ in *P. compressa* were unsuccessful due to insufficient loading of SNARF-1 in the cells. Increasing the concentration of SNARF1-AM during dye loading and adding additional wash steps of the isolated cells in autoclaved FSW to remove any free esterases from the cell suspension prior to dye loading did not improve the fluorescence signal in *P. compressa* cells.

### Statistical analyses

All analyses were conducted in R software v3.6.0 (R Core Team, 2020). Differences in reef-wide bleaching at the peak of the bleaching response, quantified as the bleaching index, between the 2015 and 2019 events and for each species between the two events were tested using unpaired two-sample t-tests. Homogeneity of variance and normality were assessed using residual plots for all analyses. Data were also tested for normality and homogeneity through Shapiro-Wilk and F-tests, respectively. The effects of prior bleaching susceptibility (B, 2 levels: susceptible and resistant) and time (T, 4 levels: 16 September 2019, 2 October 2019, 16 October 2019, 30 October 2019) were tested for coral bleaching scores and physiological parameters using linear mixed effects models (*lme4 package*; Bates et al. 2015), with colony pair included as a random intercept. The significance of fixed effects and their interactions were determined using a Type III ANOVA. Significant interactive effects were followed by pairwise comparison of estimated marginal means using Tukey adjusted p values (emmeans package, Lenth et al. 2020). Physiological measurements (gross photosynthesis, respiration, alpha, I_k_, dark-adapted yield, biomass, symbiont density, total protein content, chlorophyll content) were log-transformed to meet the assumptions of normality. To analyze the pH_i_ time series, ambient pH_i_ setpoint, acidification magnitude, and pH_i_ recovery rate, we used linear mixed effects models with bleaching susceptibility (B), cell type (C), and time (T) as fixed effects and colony pair as a random intercept. The significance of fixed effects and their interactions were determined using a Type III ANOVA.

## Results

### Lower reef-wide bleaching extent in 2019 driven by species-specific differences in bleaching response between repeat bleaching events

A marine heatwave in summer 2019 resulted in sustained temperatures above the local bleaching threshold (Fig. 1A). Accumulated heat stress peaked at 3.27 DHW in July and was sustained through the end of November (Fig. 1A). This was below that of the previous marine heatwave in 2015, which peaked at 5.84 DHW, yet was shorter in duration than 2019 (July-October; Fig. 1A). Both heatwaves led to widespread coral bleaching, although the extent and severity of bleaching across the reef was lower in 2019 than 2015 (Fig. 1B). During both events, most corals had visually recovered by December (Fig. 1B). The two dominant reef-building species in Kāne’ohe Bay, *P. compressa* and *M. capitata*, showed contrasting responses to the two heatwaves. In 2015, the maximum bleaching index was greater for *P. compressa* than *M. capitata* (Fig. 1C-D), whereas the opposite pattern was observed in 2019 (Fig. 1C-D). This was indicated by a large decrease in the bleaching index for *P. compressa* during the 2019 bleaching event relative to 2015 (p = 0.0004, t-test; Fig. 1D), but no significant difference between the two bleaching events for *M. capitata* (p = 0.7727, t-test; Fig. 1C).

### Relative bleaching susceptibility was consistent within but not between species during repeat bleaching events

Near the beginning of the 2019 heatwave in July, bleaching susceptible *M. capitata* individuals showed signs of moderate bleaching, whereas individuals with a history of bleaching resistance appeared fully pigmented (Fig. 2A-C). At the peak of heatwave in early October, bleaching susceptible *M. capitata* exhibited severe bleaching while individuals with a history of bleaching resistance remained pigmented (Fig. 2B,C; p_BxT_<0.0001, Table 1). Similarly, bleaching resistant *P. compressa* individuals remained pigmented throughout the 2019 heatwave (Fig. 2D-F; Table 1), while bleaching susceptible individuals showed significantly more paling (Fig. 2F; p_B_=0.0231, p_T_=0.0279, Table 1).

**Table 1.** Results of statistical analyses of coral bleaching scores from July 2019 - January 2020 using linear mixed effect models. Random intercept= pair. Model class = lmer. df = degrees of freedom. Significant effects determined by Chi Squared Type III ANOVA (p<0.05) are indicated in bold.

| Response Variable | Subset Analysis | Fixed Effect | Chi Squared | df | *p* |
| --- | --- | --- | --- | --- | --- |
| Bleaching Score | *Montipora capitata* | **Bleaching** | **205.444** | **1** | **< 0.0001** |
|  |  | **Time** | **90.222** | **5** | **< 0.0001** |
|  |  | **Bleaching x Time** | **45.889** | **5** | **< 0.0001** |
|  | *Porites compressa* | **Bleaching** | **5.1636** | **1** | **0.0231** |
|  |  | **Time** | **12.5583** | **5** | **0.0279** |
|  |  | Bleaching x Time | 2.4862 | 5 | 0.7785 |

### Performance of bleaching susceptible vs. bleaching resistant corals

Photosynthetic rates for *M. capitata* were lower in bleaching susceptible corals, but experienced a steep decline in both bleaching susceptible (−56.7%) and bleaching resistant corals (−63.3%) relative to mid-September, followed by a partial recovery in late October (p_T_<0.0001; p_B_=0.0202; Table 2; Fig. 3A). For *P. compressa*, time but not bleaching susceptibility was a significant factor affecting photosynthetic rates (p_T_<0.0001; Table 2), which, similar to *M. capitata*, also showed a large decline (−43% susceptible, -50.5% resistant) at the beginning of October (Fig. 3B). The initial slope of the PI curve (alpha) declined at peak bleaching for all corals, yet was steeper for resistant colonies of both *M. capitata* (p_T_<0.0001; p_B_<0.001; Table 2; Fig. 3C; Fig. S1) and *P. compressa* (p_T_<0.0001; p_B_=0.0415; Table 2; Fig. 3D) while the light saturation point (I_k_) increased at peak bleaching for both species (*M*.*capitata* p_T_<0.0001, *P. compressa* p_T_<0.0001; Table 2; Fig. 3E,F). Photochemical yield also declined at this time point for both bleaching susceptible and bleaching resistant colonies of *M. capitata* (p_T_<0.0001; Table 2; Fig. 3G) and *P. compressa* (p_T_<0.0001; Table 2; Fig. 3H), yet was higher for bleaching resistant than susceptible *M. capitata* colonies and lower for bleaching resistant than susceptible *P. compressa* colonies. (*M. capitata* p_B_=0.0056, *P. compressa* p_B_=0.0446; Table 2). Symbiont densities were lower in bleaching susceptible than resistant *M. capitata* colonies(p_B_<0.0001; Table 2; Fig. 3I). In contrast, symbiont densities in *P. compressa* were not significantly affected by either time or bleaching susceptibility (Fig. 3J, Table 2). Chlorophyll content showed similar patterns as symbiont densities, as *M. capitata* chlorophyll content was lower in bleaching susceptible corals (Fig. 3K) and not influenced by time (p_H_<0.0001; Table 2), while for *P. compressa* chlorophyll content was not significantly affected by either time or bleaching susceptibility (Fig. 3L, Table 2).

**Table 2.** Results of statistical analyses of coral performance from September - October 2019 using linear mixed effect models. All data were log10-transformed. Random intercept = pair. Model class = lmer. df = degrees of freedom (Num, Den). Significant effects determined by Satterthwaite’s Type III ANOVA (p<0.05) are indicated in bold.

| Response Variable | Subset Analysis | Fixed Effect | SS | df | F | *p* |
| --- | --- | --- | --- | --- | --- | --- |
| Gross Photosynthesis<br>($\mu\text{mol O}_2 \text{ cm}^{-2} \text{ hr}^{-1}$) | *Montipora capitata* | **Bleaching** | **0.11329** | **1, 63** | **5.6796** | **0.0202** |
|  |  | **Time** | **1.71062** | **3, 63** | **28.5869** | **< 0.0001** |
|  |  | Bleaching x Time | 0.10467 | 3, 63 | 1.7491 | 0.1661 |
|  | *Porites compressa* | Bleaching | 0.05590 | 1, 63 | 2.9911 | 0.0886 |
|  |  | **Time** | **0.78004** | **3, 63** | **13.9128** | **< 0.0001** |
|  |  | Bleaching x Time | 0.05626 | 3, 63 | 1.0034 | 0.3973 |
| Respiration<br>($\mu\text{mol O}_2 \text{ cm}^{-2} \text{ hr}^{-1}$) | *Montipora capitata* | **Bleaching** | **0.3202** | **1, 63** | **7.6755** | **0.0073** |
|  |  | **Time** | **3.5962** | **3, 63** | **28.7356** | **< 0.0001** |
|  |  | Bleaching x Time | 0.3001 | 3, 63 | 2.3977 | 0.0763 |
|  | *Porites compressa* | Bleaching | 0.00097 | 1, 63 | 0.0294 | 0.8645 |
|  |  | **Time** | **2.34608** | **3, 63** | **23.7152** | **< 0.0001** |
|  |  | Bleaching x Time | 0.13447 | 3, 63 | 1.3593 | 0.2634 |
| Alpha | *Montipora capitata* | **Bleaching** | **0.99175** | **1, 63** | **18.3635** | **< 0.0001** |
|  |  | **Time** | **1.69871** | **3, 63** | **10.4846** | **< 0.0001** |
|  |  | Bleaching x Time | 0.03932 | 3, 63 | 0.2427 | 0.8662 |
|  | *Porites compressa* | **Bleaching** | **0.15325** | **1, 63** | **4.3299** | **0.0415** |
|  |  | **Time** | **2.04907** | **3, 63** | **19.2973** | **< 0.0001** |
|  |  | Bleaching x Time | 0.03029 | 3, 63 | 0.2852 | 0.8359 |
| $I_K$ | *Montipora capitata* | Bleaching | 0.05662 | 1, 63 | 3.9361 | 0.0511 |
|  |  | **Time** | **0.39408** | **3, 63** | **9.1315** | **< 0.0001** |
|  |  | Bleaching x Time | 0.02104 | 3, 63 | 0.4874 | 0.6921 |
|  | *Porites compressa* | Bleaching<br>**Time**<br>Bleaching x Time | 0.03266<br>**0.53805**<br>0.01614 | 1, 63<br>**3, 63**<br>3, 63 | 2.5752<br>**14.1436**<br>0.4244 | 0.1136<br>**< 0.0001**<br>0.7362 |
| Dark-Adapted Yield<br>($F_v/F_m$) | *Montipora capitata* | Bleaching<br>**Time**<br>Bleaching x Time | **0.00828**<br>**0.04108**<br>0.00196 | **1, 63**<br>**3, 63**<br>3, 63 | **8.2413**<br>**13.6201**<br>0.6511 | **0.0056**<br>**< 0.0001**<br>0.5852 |
|  | *Porites compressa* | Bleaching<br>**Time**<br>Bleaching x Time | **0.00633**<br>**0.09174**<br>0.00636 | **1, 63**<br>**3, 63**<br>3, 63 | **4.2015**<br>**20.2987**<br>1.4071 | **0.0446**<br>**< 0.0001**<br>0.2490 |
| Biomass<br>($\text{mg cm}^{-2}$) | *Montipora capitata* | Bleaching<br>**Time**<br>Bleaching x Time | 0.02243<br>**0.71247**<br>0.03662 | 1, 63<br>**3, 63**<br>3, 63 | 0.7988<br>**8.4591**<br>0.4348 | 0.3749<br>**< 0.0001**<br>0.7288 |
|  | *Porites compressa* | Bleaching<br>Time<br>Bleaching x Time | 0.05549<br>0.01920<br>0.00825 | 1, 63<br>3, 63<br>3, 63 | 1.8009<br>0.2077<br>0.0892 | 0.1844<br>0.8907<br>0.9657 |
| Symbiont Density<br>($\text{cells cm}^{-2}$) | *Montipora capitata* | Bleaching<br>Time<br>Bleaching x Time | **2.40553**<br>0.22905<br>0.66737 | **1, 63**<br>3, 63<br>3, 63 | **20.1717**<br>0.6402<br>1.8654 | **< 0.0001**<br>0.5920<br>0.1445 |
|  | *Porites compressa* | Bleaching<br>Time<br>Bleaching x Time | 0.08444<br>0.41267<br>0.07025 | 1, 63<br>3, 63<br>3, 63 | 0.5196<br>0.8465<br>0.1441 | 0.4737<br>0.4736<br>0.9331 |
| Total Protein Content<br>($\text{mg cm}^{-2}$) | *Montipora capitata* | Bleaching<br>**Time**<br>Bleaching x Time | **0.19552**<br>**0.33075**<br>0.05125 | **1, 63**<br>**3, 63**<br>3, 63 | **9.9098**<br>**5.5878**<br>0.8658 | **< 0.0001**<br>**< 0.0001**<br>0.4636 |
|  | *Porites compressa* | Bleaching<br>Time<br>Bleaching x Time | 0.00579<br>0.00603<br>0.04016 | 1, 63<br>3, 63<br>3, 63 | 0.2456<br>0.0853<br>0.5794 | 0.6217<br>0.9679<br>0.6304 |
| Chlorophyll Content<br>($\mu\text{g cm}^{-2}$) | *Montipora capitata* | Bleaching<br>Time<br>Bleaching x Time | **0.91559**<br>0.71081<br>0.65168 | **1, 63**<br>3, 63<br>3, 63 | **7.5036**<br>1.9418<br>1.7802 | **< 0.0001**<br>0.1319<br>0.1600 |
|  | *Porites compressa* | Bleaching<br>Time<br>Bleaching x Time | 0.00579<br>0.00603<br>0.04101 | 1, 63<br>3, 63<br>3, 63 | 0.2456<br>0.0853<br>0.5794 | 0.6217<br>0.9679<br>0.6304 |

**Figure 3.**
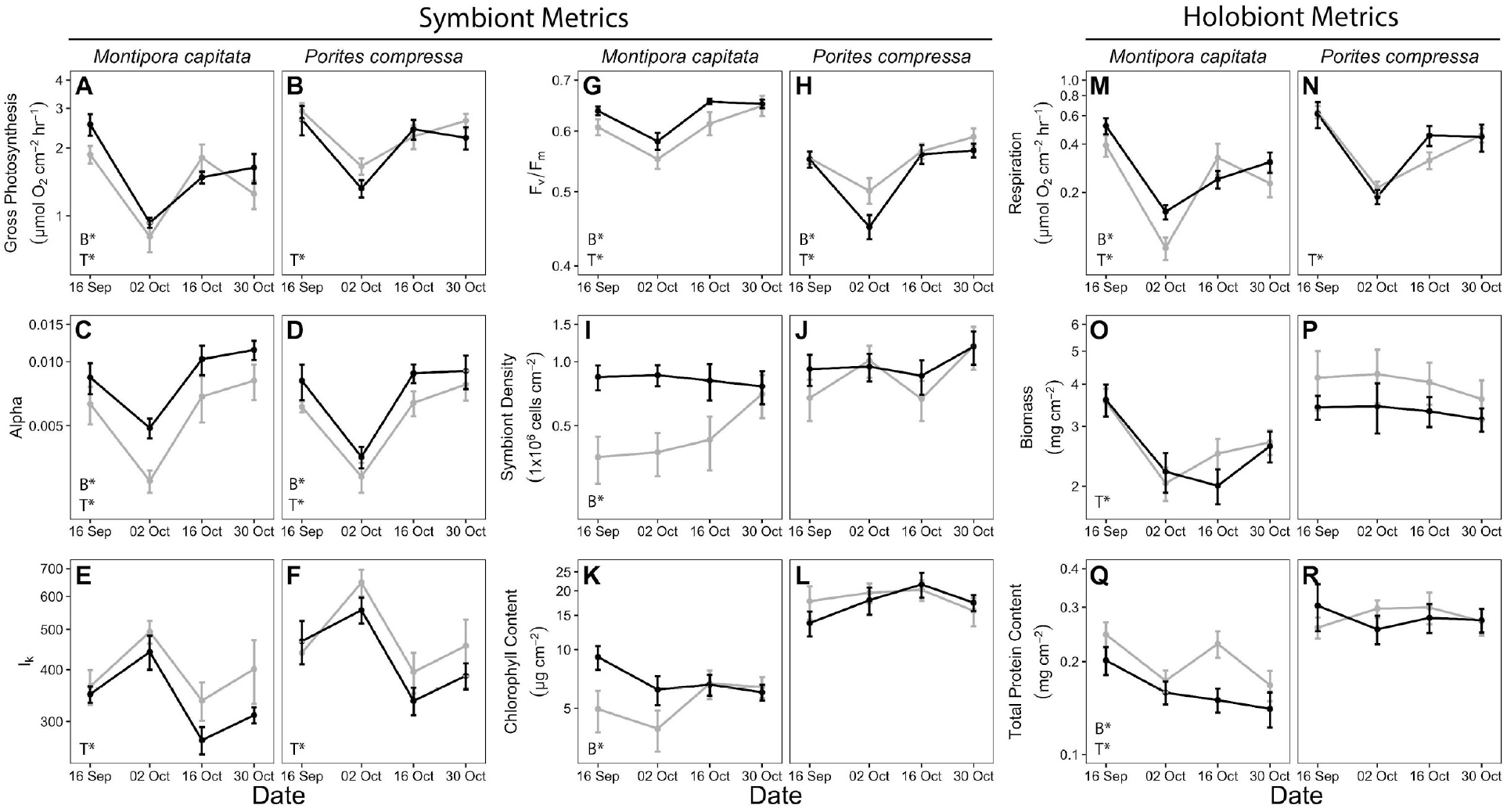
Physiological responses of bleaching susceptible (gray lines) and bleaching resistant (black lines) colonies of *Montipora capitata* and *Porites compressa* during a repeat heat stress event in 2019. A-B) gross photosynthetic rate (Pmax - LEDR); C-D) alpha (initial slope); E-F) I_k_ (light saturation point); G-H) photosynthetic efficiency (dark adapted yield; Fv/Fm); I-J) symbiont cell density; K-L) chlorophyll content; M-N) light enhanced dark respiration rate (LEDR); O-P) tissue biomass (AFDW) and Q-R) total protein content. N=10; error bars indicate SEM. Insets indicate statistical significance of bleaching susceptibility (B*) or time (T*) as determined from linear mixed effects models (p<0.05). Interactions between fixed effects were not significant for any of the metrics.

Similar to symbiont photosynthetic rates, *M. capitata* light enhanced dark respiration rates (LEDR) declined from September to early October (−76.9% susceptible, -70.8% resistant) and were generally lower in bleaching susceptible than resistant corals (p_T_<0.0001, p_B_=0.0073; Table 2; Fig. 3M). *P. compressa* LEDR experienced a similar decline (−66.2% susceptible, - 69.4% resistant) in early October, and was only affected by time (p_T_<0.0001; Table 2; Fig. 3N). Holobiont biomass was not affected by bleaching susceptibility for either species (Fig. 3O,P), but *M. capitata* biomass declined from mid-September to early October (−42.5% susceptible, -38.6% resistant) before slight recovery in late October (p_T_<0.0001; Table 2; Fig. 3O). In contrast, *P. compressa* biomass did not change through time (Fig. 3P). Total protein content decreased over time for both phenotypes and was lower in bleaching resistant than susceptible *M. capitata* (p_T_<0.0001, p_B_<0.0001; Table 2; Fig. 3Q), but was unaffected by either time or bleaching susceptibility for *P. compressa* (Table 2; Fig. 3R). Total antioxidant capacity was unaffected by time or bleaching susceptibility for both *M. capitata* (Fig. S2K) and *P. compressa* (Fig. S2L; Table S1).

### Coral acid-base homeostasis affected by both bleaching status and heat stress

Coral cells isolated from *M. capitata* (e.g. Fig. 4A) exhibited lower intracellular pH (pHi) setpoints in non-symbiocytes than symbiocytes (p_C_<0.0001) and lower pHi setpoints in cells from bleaching susceptible colonies than cells of the same type from bleaching resistant colonies (p_B_<0.0001, Table 3; Fig. 4B; Fig. S3A). When isolated coral cells were exposed to low pH seawater (pH_SW_ 7.4), pHi decreased during the first 5 min of exposure for both symbiocytes and non-symbiocytes from colonies of both bleaching histories (Fig. 4B). The magnitude of cytosolic acidification (i.e. acidosis) was greater for symbiocytes than non-symbiocytes, although bleaching susceptibility did not influence this response (p_C_ = 0.0068; Table 3; Fig. 4B; Fig. S3B). After the initial acidification, pHi of both cell types and bleaching phenotypes remained steady (Fig. 4B), with a pHi recovery rate near zero for the following 65 min that did not differ between the cell types or bleaching susceptibilities (Table 3; Fig. 4B; Fig. S3C).

**Figure 4.**
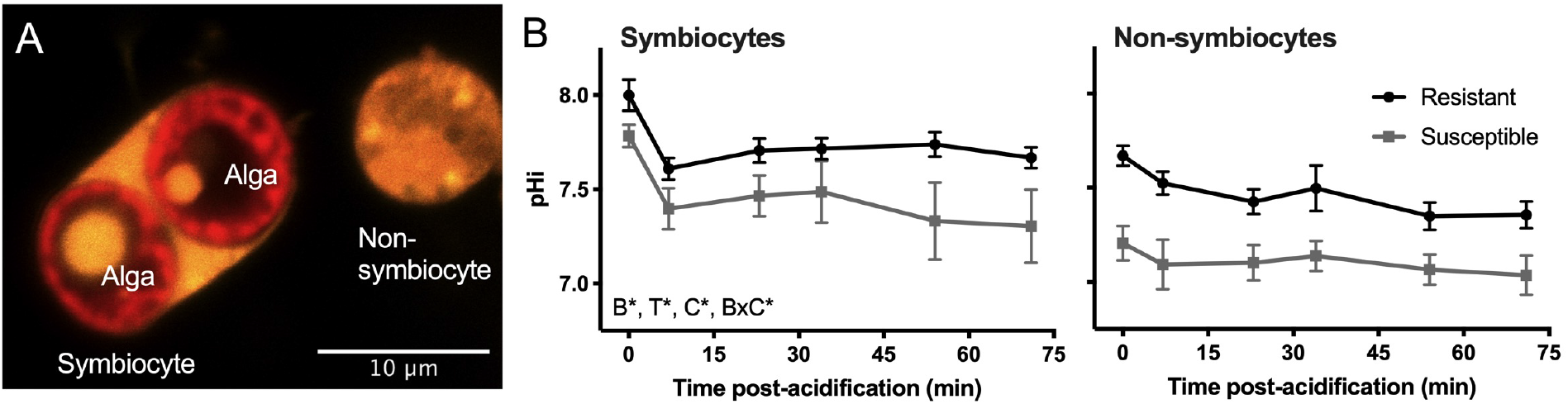
Effects of heat stress and bleaching on cellular acid-base homeostasis of *Montipora capitata*. A) Fluorescence micrograph of *M. capitata* cells containing algal symbionts (symbiocytes) and lacking algal symbionts (non-symbiocytes) loaded with SNARF1 (orange). Symbiont autofluorescence is shown in red (Alga). B) Intracellular pH (pH_i_) of coral cells following exposure to acidified seawater (pH 7.4) for symbiocytes (left) and non-symbiocytes (right). At least 8-10 cells of each cell type were analyzed per time point from each of the 10 colonies of each phenotype. Error bars indicate SEM. Letters indicate significant factors from a mixed effects model (p<0.05; B, bleaching susceptibility; C, cell type; T, time).

**Table 3.**
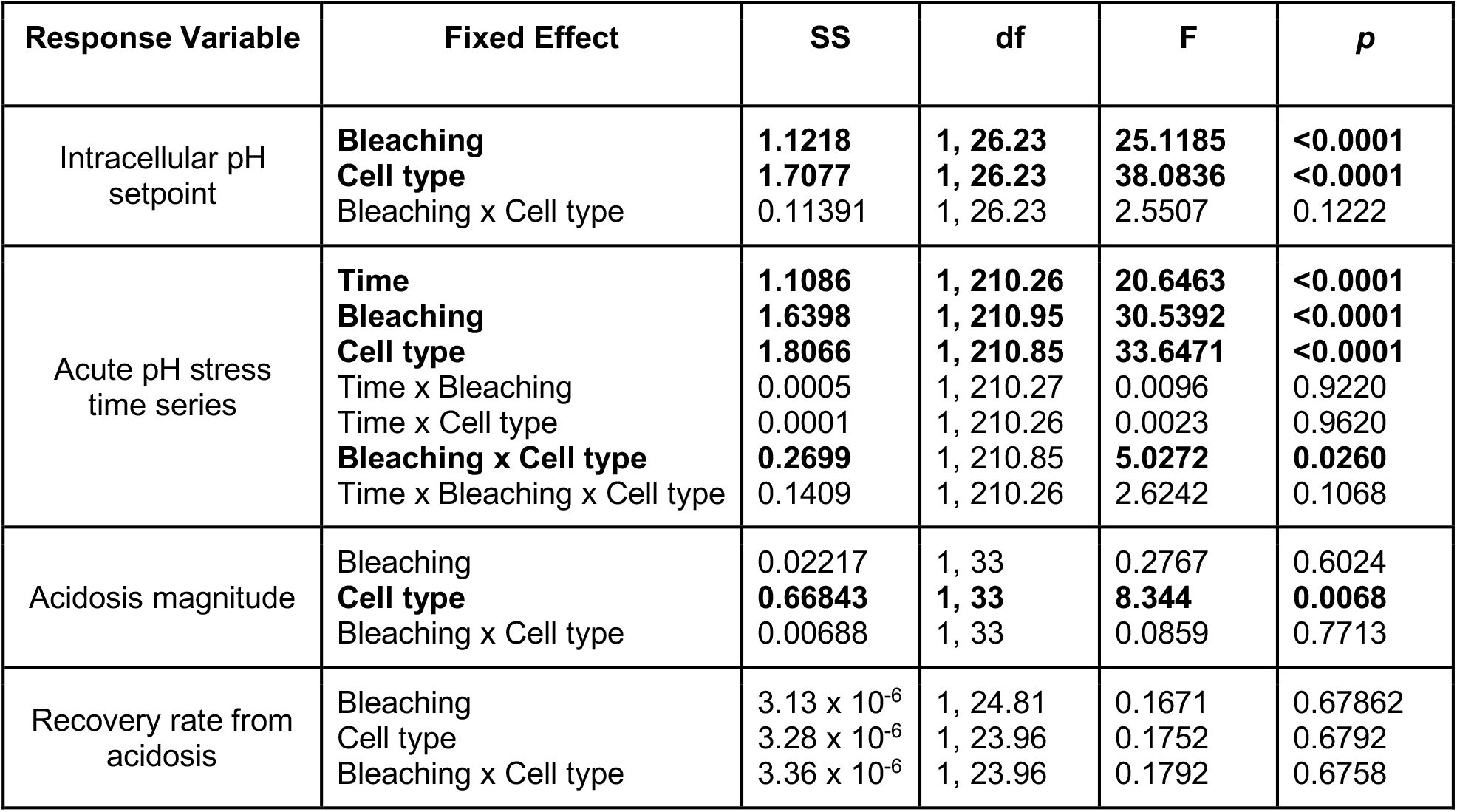
Results of statistical analyses of *Montipora capitata* host intracellular pH dynamics in response to exposure to seawater pH 7.4. Random intercept = pair. Model class = lmer. df = degrees of freedom (Num, Den). Significant effects determined by Satterthwaite’s III ANOVA (p<0.05) are indicated in bold.

| Response Variable | Fixed Effect | SS | df | F | *p* |
| --- | --- | --- | --- | --- | --- |
| Intracellular pH setpoint | **Bleaching**<br>**Cell type**<br>Bleaching x Cell type | **1.1218**<br>**1.7077**<br>0.11391 | **1, 26.23**<br>**1, 26.23**<br>1, 26.23 | **25.1185**<br>**38.0836**<br>2.5507 | **<0.0001**<br>**<0.0001**<br>0.1222 |
| Acute pH stress time series | **Time**<br>**Bleaching**<br>**Cell type**<br>Time x Bleaching<br>Time x Cell type<br>**Bleaching x Cell type**<br>Time x Bleaching x Cell type | **1.1086**<br>**1.6398**<br>**1.8066**<br>0.0005<br>0.0001<br>**0.2699**<br>0.1409 | **1, 210.26**<br>**1, 210.95**<br>**1, 210.85**<br>1, 210.27<br>1, 210.26<br>1, 210.85<br>1, 210.26 | **20.6463**<br>**30.5392**<br>**33.6471**<br>0.0096<br>0.0023<br>**5.0272**<br>2.6242 | **<0.0001**<br>**<0.0001**<br>**<0.0001**<br>0.9220<br>0.9620<br>**0.0260**<br>0.1068 |
| Acidosis magnitude | Bleaching<br>**Cell type**<br>Bleaching x Cell type | 0.02217<br>**0.66843**<br>0.00688 | 1, 33<br>**1, 33**<br>1, 33 | 0.2767<br>**8.344**<br>0.0859 | 0.6024<br>**0.0068**<br>0.7713 |
| Recovery rate from acidosis | Bleaching<br>Cell type<br>Bleaching x Cell type | $3.13 \times 10^{-6}$<br>$3.28 \times 10^{-6}$<br>$3.36 \times 10^{-6}$ | 1, 24.81<br>1, 23.96<br>1, 23.96 | 0.1671<br>0.1752<br>0.1792 | 0.67862<br>0.6792<br>0.6758 |

## Discussion

### Changes in relative bleaching susceptibility between species during repeat heatwaves indicate possible trade-offs between coral bleaching resistance vs. resilience

Here we found that relative bleaching susceptibility was consistent within species between two bleaching events separated by four years, indicating that these species appear to have fixed differences in relative bleaching susceptibility between individuals of the same species. Patterns of differential bleaching susceptibility are consistent with the responses of these same two species to a previous repeat bleaching event (2014 and 2015; Ritson-Williams and Gates 2020) and during a simulated marine heatwave (Dilworth et al. 2021). Surprisingly, bleaching susceptible individuals of each species in our study responded differently to the second heatwave (2019), indicating possible differences in acclimatization and recovery dynamics. For example, bleaching susceptible *M. capitata* bleached severely in both events, despite lower levels of heat stress in 2019 than in 2015, indicating that this species did not show acclimatization. In contrast, bleaching susceptible *P. compressa* exhibited severe bleaching in 2015 but only mild bleaching in 2019, perhaps corresponding with the lower levels of heat stress in the second event, or as a result of acclimatization during the first event, making them more resistant to heat stress upon repeat exposure. That *P. compressa* performed better than *M. capitata* in 2019 was unexpected given that in 2015, *P. compressa* had greater susceptibility to bleaching (both greater severity and prevalence) than *M. capitata* across the population (Matsuda et al. 2020). However, bleached *P. compressa* did recover more quickly than *M. capitata* after the 2015 event, and this rapid recovery in *P. compressa* corresponded with lower levels of partial mortality in bleaching susceptible *P. compressa* than *M. capitata* in the two years following the heatwave (Matsuda et al. 2020). This is somewhat counterintuitive, as bleaching itself is often predictive of mortality, and bleaching susceptible species tend to be the first to be lost from the population (Hughes et al. 2018; McClanahan 2004; Loya et al. 2001; Eakin et al. 2010).

These differences in bleaching recovery dynamics following the 2015 event may help explain the differences in bleaching severity observed during the second heatwave. In the context of repeat bleaching, the rapid recovery and lower partial mortality of bleaching susceptible *P. compressa* likely meant that these individuals were in better energetic standing than bleaching susceptible *M. capitata* going into the 2019 event. In particular, the slow recovery rates of these same pairs of *M. capitata* (Matsuda et al 2020) may have left them energetically depleted and thus hypersensitive to severe bleaching during the repeat heatwave, despite lower levels of heat stress in the second event. Indeed, even four years following the 2015 bleaching event, bleaching susceptible colonies of *M. capitata* had distinct lipid profiles from bleaching resistant colonies despite both phenotypes appearing visually healthy (Roach et al. 2021), supporting the hypothesis that recovery from that event prior to the 2019 event was incomplete in this species. A slow energetic recovery has been shown to make individuals less resistant to bleaching and mortality following repeat exposure to heat stress in other coral species (Sale et al. 2019), highlighting a barrier to acclimatization and the uneven harm of repeat bleaching on different coral species. The tradeoff with greater bleaching resistance in *M. capitata* for slower recovery from bleaching indicates that this strategy is only advantageous when heat stress events are far enough apart for complete recovery to occur. The downsides of this tradeoff are apparent in other species when heat stress is particularly prolonged, as corals initially more resistant to bleaching no longer had a survival advantage when heat stress was sustained for an unprecedented 10 months (Claar et al. 2020). This would indicate that the duration and magnitude of heat stress, as well as that of the subsequent recovery periods, control the tradeoff between resistance and recovery, and may be reversing the relative advantages between species that occur during isolated heat stress events. This has been observed in other reef systems as well (Grottoli et al. 2014), indicating this pattern may be common across reef systems.

### Metabolic depression occurred in heat stressed corals regardless of bleaching

All corals examined here exhibited metabolic depression and a decline in photochemical capacity during the 2019 marine heatwave irrespective of bleaching phenotype. This decline was temporary, as metabolic rates for both phenotypes began recovering as soon as 2 weeks following peak metabolic stress. Metabolic recovery lagged behind symbiont repopulation in bleaching susceptible individuals, and the decline persisted in both phenotypes for several weeks after temperatures dropped below the bleaching threshold. For bleaching susceptible *P. compressa*, which suffered only mild paling, the decline in metabolic performance did not result in differences in overall tissue biomass or protein content, indicating that these physiological effects were temporary enough that they did not manifest in significant energetic differences in tissue reserves. In contrast, bleaching susceptible *M. capitata* had lower symbiont abundance and chlorophyll content than bleaching resistant corals, but both phenotypes exhibited declines in tissue biomass over time. These responses indicate that bleaching resistant corals were still stressed even though their symbionts remained, and that both phenotypes may have switched to meet their energetic demands by the catabolism of energy-rich biomass (i.e., proteins, lipids, and carbohydrates; Fitt et al. 1993; Grottoli, Rodrigues, and Palardy 2006; Schoepf et al. 2015; Rädecker et al. 2021). It is also possible that heat stress led to a reduction in carbon translocation from the symbiont to the host, as others have observed a switch in metabolic exchanges between host and symbiont during heat stress that occurs prior to bleaching (Rädecker et al. 2021; Baker et al. 2018). Differences in symbiont loss and biomass reductions in resistant and susceptible *M. capitata* could also have been due to differences in symbiont species hosted, as bleaching resistant *M. capitata* colonies in this study primarily hosted *Durusdinium* sp. with some *Cladocopium* sp, as opposed to the bleaching susceptible individuals, which hosted almost exclusively *Cladocopium* sp. (Dilworth et al. 2021). *Durusdinium*, while generally more heat tolerant (Stat and Gates 2010), are often less generous symbionts (e.g. Cunning et al. 2015); however, whether there is a tradeoff with heat tolerance and symbiont generosity or a relationship between generosity and temperature for *Durusdinium* spp. associated with *M. capitata* remains unknown. Furthermore, hosts like *M. capitata* with flexible symbiont associations are often more susceptible to bleaching than hosts like *P. compressa* that strictly associate with only a single symbiont species (Putnam et al. 2012), which may help explain the species specific differences in repeat heat stress responses.

The energetic deficits incurred during heat stress due to declining symbiont contributions and reallocations of energy expenditure towards maintenance may come at the cost of ATP-demanding processes like growth, reproduction, and feeding activity (Sokolova et al. 2012). While metabolic depression can extend an organism’s survival during heat stress (H. Pörtner 2008), these changes in energy allocation have long-term potential consequences for coral fitness for all individuals exposed to temperatures above historical summer averages, as even small changes in energy balance (e.g. declines in long-term energy stores like lipids or short-term stores like ATP) can cause substantial ecological disadvantages (Anthony et al. 2009; H. Pörtner 2008). In corals, the harmful effects of heat stress can persist for months (A. D. Hughes and Grottoli 2013; Baumann et al. 2014) to years (Levitan et al. 2014; Cantin and Lough 2014; Matsuda et al. 2020), even after the symbiosis is reestablished and even for corals that never bleached. The harmful effects of bleaching do compound those of heat stress alone, as bleaching susceptible corals have reduced spawning frequency and fecundity than bleaching resistant corals of the same species (Levitan et al. 2014; Fisch et al. 2019; Howells et al. 2016). However, the evident consequences of heat stress could mean that acclimatization to high temperatures that results in a decline in bleaching during heatwaves does not necessarily equate to resilience, given that heat stress substantially reduces coral growth and reproduction even in bleaching resistant individuals (Levitan et al. 2014; Fisch et al. 2019; Howells et al. 2016; Cantin and Lough 2014), reducing fitness and potentially altering the ecological dynamics of the entire ecosystem.

### Heat stress differentially alters intracellular pH and impairs cellular acidification resilience in bleaching resistant and bleaching susceptible colonies

Here we found that intracellular pH (pH_i_) setpoints were depressed in bleaching susceptible corals relative to bleaching resistant corals during heat stress, even in the absence of external pH stress. This is the first observation of differences in pH_i_ setpoints due to thermal stress in a cnidarian. Because impaired pH_i_ regulation occurred in all coral cell types, including those that still harbored symbionts, it appears that changes in symbiont abundance throughout the colony due to heat stress were sufficient to cause pH_i_ dysregulation throughout the animal. Experimental tests of intracellular acidification resilience demonstrated that the magnitude of intracellular acidosis was greater in symbiocytes than non-symbiocytes following exposure to pH stress. This was somewhat surprising, because the higher pH_i_ setpoints of symbiocytes would be expected to yield greater buffering capacity due to greater passive buffering by bicarbonate (Boron 2004).

The greater acidosis in symbiocytes following acute pH stress could therefore indicate that the presence of heat-stressed symbionts may be harmful to metabolic activity and homeostatic functions of the host cell. Furthermore, because cells from bleaching susceptible colonies started at a lower pH_i_ setpoint, the pH_i_ at maximum acidosis was lower than in cells from bleaching resistant colonies, indicating a compounding effect of colony sensitivity to heat stress on cellular pH dysregulation during acidification stress. Previous work in *M. capitata* has shown a link between heat stress and cellular acidification, as cells exposed to pH stress exhibited lower pH_i_ during acidosis as temperatures increased (Gibbin et al. 2015). Here, however, these differences were observed at the same temperature within a single species, indicating that greater intracellular acidosis following exposure to low external pH may be driven by energetic deficits due to the loss of symbionts, and perhaps energetic reserves, throughout the colony. Indeed, species-specific differences in symbiont loss during heat stress corresponded with differences in pH_i_ following exposure to low pH seawater, where greater symbiont loss led to greater acidosis. Here, we saw a similar pattern within a single species, suggesting that this response may be due to colony-specific differences and not species-specific differences in bleaching thresholds.

Heat stress also rendered coral cells unable to recover from experimentally induced acidosis. This is the first study to our knowledge to document a lack of compensatory response in corals. In the few cnidarian studies conducted to date, both symbiocytes and non-symbiocytes exposed to low pH seawater initially acidify, then return to their pH_i_ setpoint within ∼75 minutes. This occurs under both light (Gibbin et al. 2014) and dark conditions (Laurent et al. 2014; Barott, Barron, and Tresguerres 2017), indicating that corals have compensatory mechanisms that are upregulated in response to acidosis (e.g. ion exchangers; (Laurent et al. 2014)) and this response is not solely due to buffering by the symbiont. Sustained exposure to heat stress *in situ* curtailed corals’ ability to compensate for intracellular acidosis in the dark for both symbiocytes and non-symbiocytes. This pattern was consistent between cells from bleaching susceptible and resistant corals, suggesting that heat stress itself was sufficient to sensitize corals to acid stress even if they retained their symbionts.

The mechanisms by which bleaching and heat stress disrupt cellular pH dynamics remain to be described, but are likely instigated by metabolic changes in the host. For example, metabolic depression due to thermal stress in various marine vertebrates and invertebrates can cause a decrease in H^+^ excretion that results in decreases in the pH_i_ setpoint (H. Pörtner 2008; H. O. Pörtner and Bock 2000). Our observations of a lower pH_i_ setpoint in bleaching susceptible colonies than bleaching resistant colonies is consistent with corresponding observations of more severe metabolic depression in these individuals. Acid-base homeostasis is energetically expensive, and the costs of this vital maintenance process increase following stress, and so may be particularly exacerbated during coral bleaching. There is a growing understanding of coral acid-base sensing mechanisms (Barott, Barron, and Tresguerres 2017; Barott, Venn, et al. 2020) and the downstream effectors involved in maintaining acid-base homeostasis in the cytosol (Laurent et al. 2014), calcifying fluid (Zoccola et al. 2015; Barott, Perez, et al. 2015; Zoccola et al. 2004), and symbiosome (Barott, Venn, et al. 2015; Bertucci et al. 2009; Barott, Perez, et al. 2015). While previous work in corals has shown corals can upregulate active acid-base regulation processes under ambient temperature conditions, the relative contribution of passive (e.g. bicarbonate buffering) vs. active (ATP-dependent ion transport) acid-base regulation in determining coral pH_i_ remains an outstanding question. More work is needed to understand how these pathways respond to heat stress, their capacity for acclimatization, and adaptive mechanisms of resilience between resistant and susceptible individuals and species. Initial work has found that heat stress decreases gene expression of ion transporters (bicarbonate, H^+^) and carbonic anhydrases that interconvert bicarbonate and CO_2_ (Bernardet et al. 2019; Kenkel, Meyer, and Matz 2013), reductions that correspond with metabolic depression and declines in calcification (Bernardet et al. 2019). Downregulation of ion transport pathways during heat stress could also impair the function of the symbiosis, which is dependent on delivery of inorganic carbon to symbionts, a process that is facilitated by host proton transport (Barott, Venn, et al. 2015). Indeed, these changes could contribute to the declines in photosynthesis observed in corals that retained their symbionts throughout the heatwave, and is consistent with disruptions of nutrient cycling within the symbiosis during heat stress prior to bleaching (Rädecker et al. 2021).

The sensitivity of acid-base homeostasis to heat stress poses a distinct challenge for corals, as both the symbiosis and calcification are sensitive to the acid-base status of the animal (Barott et al. 2015; Barott et al. 2020; McCulloch et al. 2012; Venn et al. 2012). Furthermore, the nature of the interactions between temperature stress and acid-base homeostasis are important for corals coping with climate change, as the associated increases in CO_2_ dissolution in the ocean may lead to hypercapnia within coral tissues, which itself can alter thermal stress responses in marine ectotherms by narrowing the range of temperatures they can tolerate (H. Pörtner 2008). This interaction may therefore increase the potential for ocean acidification to compound the harm of temperature stress on corals and exacerbate bleaching (Dove et al. 2020; Gibbin et al. 2015), especially because these animals are already living at the upper edge of their thermal tolerance windows. Furthermore, if pH_i_ in corals is regulated more tightly than extracellular body fluids (pH_e_) during stress, as it is in other marine invertebrates (Tresguerres et al. 2020), the interactive effects of temperature stress and hypercapnia on pH_i_ and pH_e_ regulation may be particularly detrimental for biomineralization, which occurs in extracellular pockets of fluid located between the coral epidermis and the skeleton. Indeed, recent work has found that heat stress impairs coral regulation of calcifying fluid pH_e_ with concurrent declines in calcification (Guillermic et al. 2021; Schoepf et al. 2021), which raises the question of whether maintenance of coral pH_i_ may be occurring at the expense of pH_e_ during stress. Because the dynamics and mechanisms of acid-base regulation differ greatly between species and cell types (Tresguerres et al. 2020, 2017), it is critical that future work describe the molecular mechanisms that dictate these organismal responses to climate change stressors in order to better predict their ability to acclimatize and adapt to the ongoing climate crisis.

### Conclusions

It is important to understand physiological processes underlying coral heat stress responses, both independent of and during bleaching, in order to better predict how corals will respond to global climate change. Here we uncovered cellular and organismal costs to *in situ* heat stress that were suffered by both bleaching susceptible and bleaching resistant corals. Thermal stress coupled with bleaching exacerbated metabolic stress at the organismal and cellular scales beyond that of heat stress alone. These results are crucially important, as disruptions in energy balance following heat stress can persist for years even if the symbiosis remains intact. Accounting for these changes is important to avoid underestimating the harmful effects of sub-bleaching heat anomalies on ecosystem function and the challenges to coral persistence in a changing ocean.

## Acknowledgements

We thank Chris Suchocki and Rayna McClintock for help with coral collections.

## Author contributions

Designed the study: KLB, HMP, CEN

Collected and analyzed the data: TI, LAW, WS, KTB, CC, EK, AH, KLB, CEN

Wrote the paper: KLB, TI, LAW, KTB

All authors edited and approved the final manuscript.

## Data accessibility

The datasets analyzed for this study and R scripts can be found in GitHub: https://github.com/BarottLab/KBayBleach2019. The raw data supporting the conclusions of this manuscript will be made available by the authors, without undue reservation, to any qualified researcher.

