## Supplemental materials for "Marine heatwaves depress metabolic activity and impair cellular acid-base homeostasis in reef-building corals regardless of bleaching susceptibility"

### Supplemental Figures

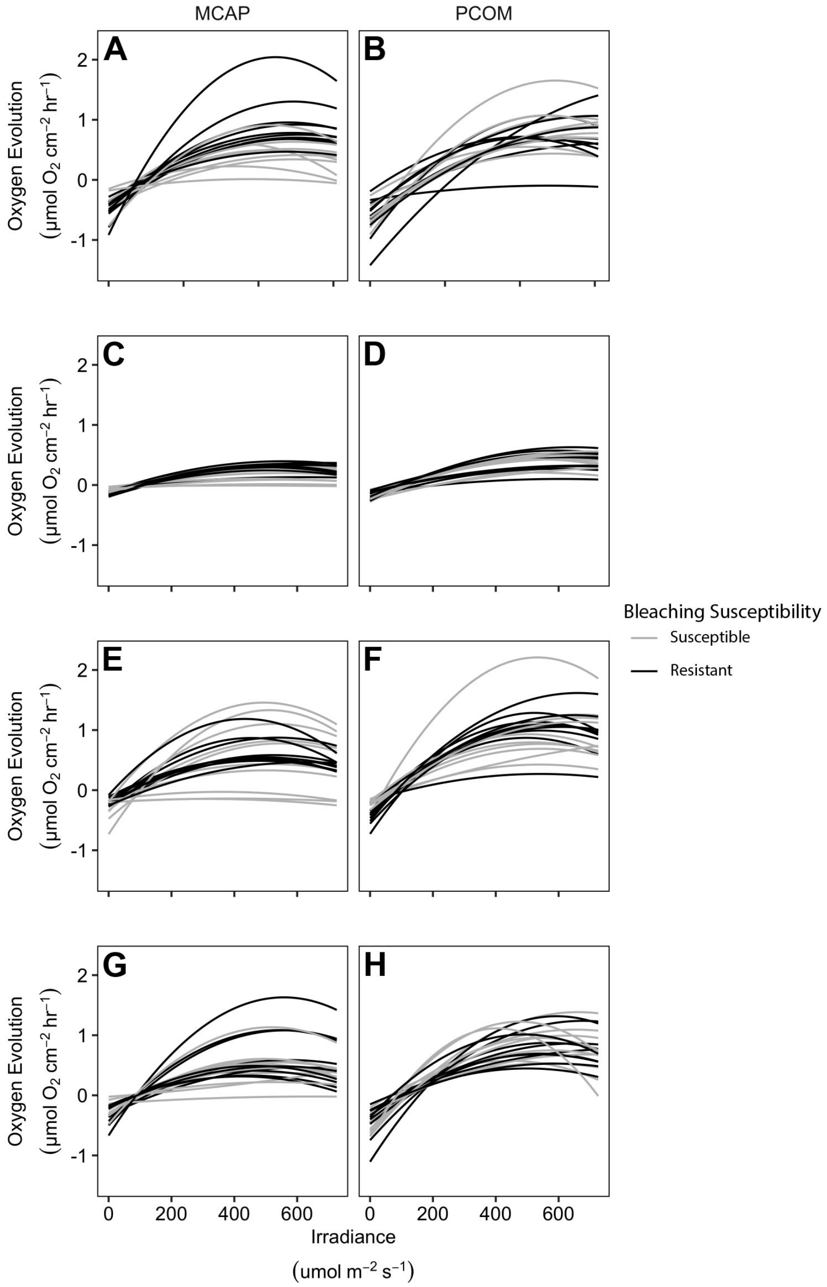


**Figure S1.** Photosynthesis-irradiance curves of bleaching susceptible and bleaching resistant colonies of *Montipora capitata* and *Porites compressa* generated by curve fitting of oxygen consumption and production rates on A-B) September 16 2019; C-D) October 2, 2019; E-F) October 16, 2019; G-H) October 30, 2019.

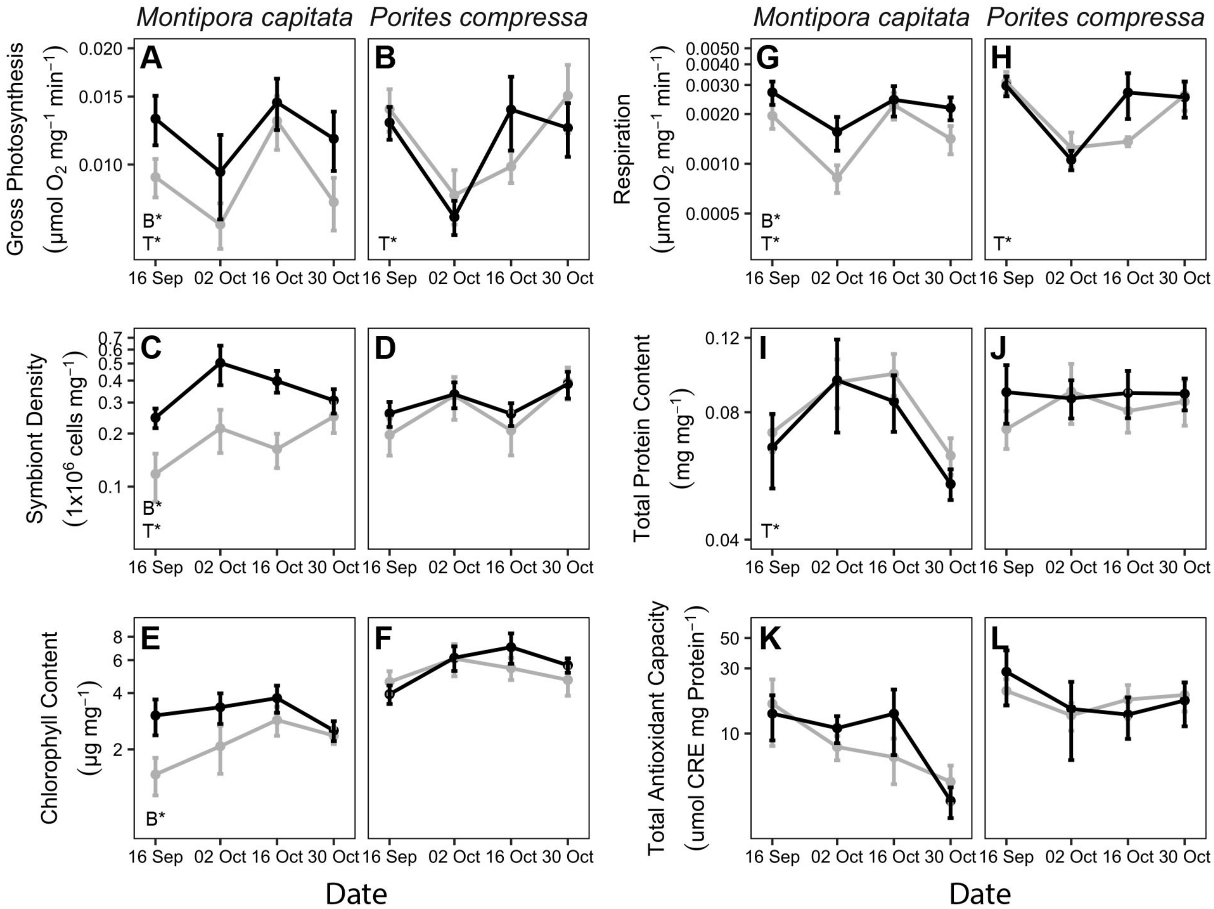


**Figure S2.** Physiological responses normalized to biomass (AFDW) of bleaching susceptible (gray lines) and bleaching resistant (black lines) colonies of *Montipora capitata* and *Porites compressa* during a repeat heat stress event in 2019. A-B) gross photosynthetic rate (Pmax - LEDR); C-D) symbiont cell density; E-F) chlorophyll content; G-H) light enhanced dark respiration rate (LEDR); I-J) total protein content; K-L) total antioxidant capacity. N=10; error bars indicate SEM. Insets indicate statistical significance of bleaching susceptibility (B*) or time (T*) as determined from linear mixed effects models ($p<0.05$).

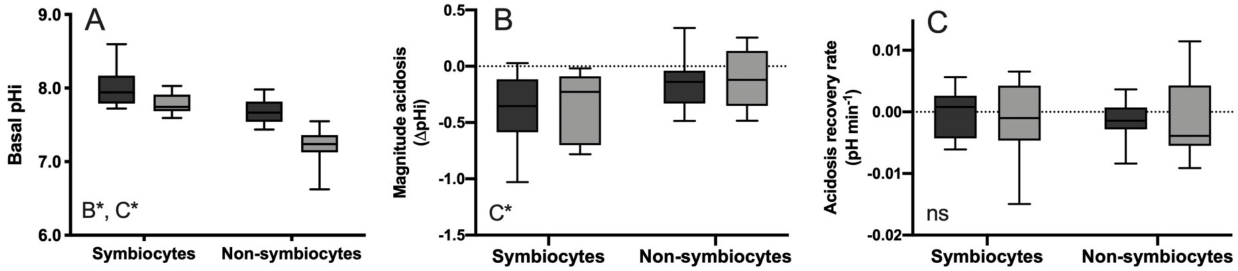


**Figure S3.** Effects of heat stress and bleaching on cellular acid-base homeostasis of *Montipora capitata*. A) Basal pHi setpoint of coral cells from bleaching resistant (dark gray) or bleaching susceptible (light gray) colonies under ambient seawater pH conditions (pH 8.0); inset indicates significant factors of bleaching susceptibility (B*) and cell type (C*) from linear mixed effect model ($p<0.0001$, 2-way ANOVA). B) Magnitude of acidosis of coral cells following 5 minutes of exposure to acidified seawater. Inset indicates significant factors ($p<0.05$, 2-way ANOVA); C) Recovery rate of pHi during the 65 minutes following maximum acidosis. No significant differences were observed (ns, $p>0.05$, 2-way ANOVA).

**Supplemental Tables**

**Table S1.** Results of statistical analyses of coral performance normalized to biomass (AFDW except where indicated) from September - October 2019 using linear mixed effect models. All data were log10-transformed. Random intercept = pair. Model class = lmer. df = degrees of freedom (Num, Den). Significant effects determined by Satterthwaite's Type III ANOVA ($p<0.05$) are indicated in bold.

| Response Variable | Subset Analysis | Fixed Effect | SS | df | F | *p* |
|---|---|---|---|---|---|---|
| Gross Photosynthesis (µmol $O_2$ $mg^{-1}$ $hr^{-1}$) | *Montipora capitata* | **Bleaching** | **0.2365** | **1, 63** | **5.0815** | **0.0272** |
| | | **Time** | **0.6260** | **3, 63** | **4.4832** | **< 0.0001** |
| | | Bleaching x Time | 0.0263 | 3, 63 | 0.1884 | 0.9040 |
| | *Porites compressa* | Bleaching | 0.0000 | 1, 63 | 0.0000 | 0.9961 |
| | | **Time** | **0.8052** | **3, 63** | **8.8656** | **< 0.0001** |
| | | Bleaching x Time | 0.0855 | 3, 63 | 0.9417 | 0.4259 |
| Respiration (µmol $O_2$ $mg^{-1}$ $hr^{-1}$) | *Montipora capitata* | **Bleaching** | **0.5120** | **1, 63** | **6.3659** | **0.0142** |
| | | **Time** | **1.5145** | **3, 63** | **6.2761** | **< 0.0001** |
| | | Bleaching x Time | 0.1377 | 3, 63 | 0.5708 | 0.6363 |
| | *Porites compressa* | Bleaching | 0.0418 | 1, 63 | 0.8276 | 0.3664 |
| | | **Time** | **2.3497** | **3, 63** | **15.5074** | **< 0.0001** |
| | | Bleaching x Time | 0.1831 | 3, 63 | 1.2085 | 0.3140 |
| Symbiont Density (cells $mg^{-1}$) | *Montipora capitata* | **Bleaching** | **2.8924** | **1, 63** | **23.1645** | **< 0.0001** |
| | | **Time** | **1.1357** | **3, 63** | **3.30320** | **0.0347** |
| | | Bleaching x Time | 0.6239 | 3, 63 | 1.6656 | 0.1820 |
| | *Porites compressa* | Bleaching | 0.2768 | 1, 63 | 1.4247 | 0.2371 |
| | | Time | 0.5831 | 3, 63 | 1.0003 | 0.3987 |
| | | Bleaching x Time | 0.0819 | 3, 63 | 0.1407 | 0.9353 |
| Total Protein (mg $mg^{-1}$) | *Montipora capitata* | Bleaching | 0.0855 | 1, 63 | 2.5888 | 0.1126 |
| | | **Time** | **0.4848** | **3, 63** | **4.8924** | **< 0.0001** |
| | | Bleaching x Time | 0.0041 | 3, 63 | 0.0415 | 0.9886 |
| | *Porites compressa* | Bleaching | 0.0254 | 1, 63 | 0.7820 | 0.3799 |
| | | Time | 0.0136 | 3, 63 | 0.1396 | 0.9359 |
| | | Bleaching x Time | 0.0192 | 3, 63 | 0.1973 | 0.8978 |
| Chlorophyll (µg $mg^{-1}$) | *Montipora capitata* | **Bleaching** | **1.2246** | **1, 63** | **8.3400** | **< 0.0001** |
| | | Time | 0.6006 | 3, 63 | 1.3634 | 0.2609 |
| | | Bleaching x Time | 0.4848 | 3, 63 | 1.1006 | 0.3546 |
| | *Porites compressa* | Bleaching | 0.0478 | 1, 63 | 0.6240 | 0.4322 |
| | | Time | 0.2443 | 3, 63 | 1.0630 | 0.3703 |
| | | Bleaching x Time | 0.1375 | 3, 63 | 0.5987 | 0.6179 |
| Total Antioxidant Capacity (µmol CRE $mg^{-1}$ protein) | *Montipora capitata* | Bleaching | 0.1202 | 1, 63 | 0.3812 | 0.5392 |
| | | Time | 2.3278 | 3, 63 | 2.4610 | 0.0708 |
| | | Bleaching x Time | 0.3744 | 3, 63 | 0.3959 | 0.7564 |
| | *Porites compressa* | Bleaching | 0.1747 | 1, 63 | 0.8292 | 0.3660 |

| | | | | | | |
|---|---|---|---|---|---|---|
| | | Time<br>Bleaching x Time | 0.6467<br>0.0672 | 3, 63<br>3, 63 | 1.0230<br>0.1063 | 0.3885<br>0.9561 |
